# Acceleration of HDL-mediated cellular cholesterol efflux alleviates periodontitis

**DOI:** 10.1101/2024.01.18.576176

**Authors:** Thanh-Tam Tran, Gyuseok Lee, Yun Hyun Huh, Ki-Ho Chung, Sun Young Lee, Ka Hyon Park, Min-Suk Kook, Jaeyoung Ryu, Ok-Su Kim, Hyun-Pil Lim, Jeong-Tae Koh, Je-Hwang Ryu

## Abstract

Periodontitis (PD) is a common inflammatory disease known to be closely associated with metabolic disorders, particularly hyperlipidemia. However, direct evidence is lacking, and the molecular mechanism is yet to be examined. In the current study, we demonstrated that hypercholesterolemia is a causative factor in the development of PD. Logistic regression analysis revealed a strong positive correlation between PD and dyslipidemia. Data from *in vivo* (PD mouse model subjected to a high cholesterol diet) and *in vitro* (cholesterol treatment of periodontal cells) experiments showed that excess cholesterol influx into periodontal cells potentially contributes to periodontal inflammation and subsequently, alveolar bone erosion. Additionally, we compared the protective efficacies of cholesterol-lowering drugs with their different modes of action against PD pathogenesis in mice. Among the cholesterol-lowering drugs we tested, fenofibrate exerted the most protective effect against PD pathogenesis, due to an increased level of high-density lipoprotein cholesterol, a lipoprotein involved in cholesterol efflux from cells and reverse cholesterol transport. Indeed, cholesterol efflux was suppressed during PD progression by downregulation of the apoA-I binding protein (APOA1BP) expression in inflamed gingival fibroblasts and periodontal ligament cells. We also demonstrated that the overexpression of APOA1BP efficiently regulated periodontal inflammation and the subsequent alveolar bone loss by inducing cholesterol efflux. Our collective findings highlight the potential utility of currently available cholesterol-lowering medications for the mitigation of PD pathogenesis. By targeting the acceleration of high-density lipoprotein (HDL)-mediated cellular cholesterol efflux, a new therapeutic approach for PD may become possible.

## INTRODUCTION

Periodontal disease is one of the most common chronic inflammatory conditions resulting in discomfort and pain^1,2^. Its prevalence ranges from 20% to 50% around the world^3^. Periodontal disease affects the gingival tissue and other structures supporting teeth, such as the alveolar bone and periodontal ligament. In the early stage, the inflammation is limited to the gingival tissue and referred to as “gingivitis”, which is characterized by bleeding, gum enlargement, and pain. Untreated gingivitis may result in the more severe form of gum disease, namely “periodontitis (PD)”, which relates to the loss of the periodontal attachment and alveolar bone destruction, causing teeth to become loose, or even fall out^2,4^.

The primary etiology of PD is a proliferation of gram-negative anaerobic species including *A. actinomycetemcomitans* and *P. gingivalis*^5,6^. However, extensive investigations have revealed that PD is not simply an inflammatory disease caused by specific pathogenic bacteria. Various potential risk factors, such as gender, smoking, alcohol, stress, diabetes, obesity and metabolic syndrome, osteoporosis, dietary calcium, and genetic factors, can significantly affect host immune responses and host susceptibility to pathogens in the oral environment^7,8^. Recently, obesity and metabolic syndrome emerged as a new research perspective for PD management because both metabolic syndrome and PD are more common in the elderly^9–12^. Previous studies have demonstrated the close association between metabolic syndrome and the development of PD, especially in terms of the lipid profile. In particular, it has been shown in patients that serum cholesterol levels positively correlate with PD ratio^13–16^. Conventional periodontal treatment options include surgical and non-surgical approaches and adjunctive therapy. These methods are generally ineffective in the presence of uncontrolled host immune responses^17^. Novel therapeutic approaches for PD require a comprehensive understanding of the individual molecular pathogenic mechanisms and efforts to develop target-oriented drugs. Since metabolic abnormalities, such as hyperlipidemia, can alter the oral microbiome and potentiate the deleterious effects of PD, improved knowledge of these associations may significantly expand the scope and appreciation of oral health and dental practices.

Cholesterol is an essential structural component of the cell membrane and serves as a substrate for the synthesis of several important biomolecules. The cellular homeostasis of cholesterol is tightly controlled through synthesis, influx, efflux, and metabolism^18^. Cholesterol efflux enables the clearance of excess cholesterol from cells and depends on the high-density lipoprotein (HDL), a complex particle comprising lipoproteins, phospholipids, and cholesterol^19^. Cholesterol homeostasis plays a central role in numerous physiological processes and is emerging as a contributory factor in several human disorders including Alzheimer’s disease and multiple sclerosis. Previous experiments by our group indicate that dysregulation of cholesterol homeostasis can result in osteoarthritis^20^. Although a few reports and various meta-analyses support a possible association between serum cholesterol and human PD, to our knowledge, no investigations have focused on whether cellular cholesterol levels are modulated in periodontal cells and their potential effects on PD pathogenesis. More importantly, whether cholesterol or its metabolites are directly associated with PD pathogenesis and the underlying mechanisms remain to be established.

This study aimed to investigate the correlations between hypercholesterolemia and PD by analyzing clinical data as well as *in vitro* and *in vivo* experimental data using regression analyses. We also investigated the protective effect of cholesterol-lowering drugs in PD, with a focus on the regulation of cholesterol efflux by HDL and APOA1BP. Our data provide direct evidence that hypercholesterolemia is a risk factor for PD and supports that the promotion of HDL-mediated cholesterol efflux can be a potential avenue for the treatment of PD as a manifestation of systemic disorder.

## RESULTS

### Accumulation of cellular cholesterol causes periodontal inflammation

To explore the association between PD pathogenesis and dyslipidemia, we analyzed the serum lipid profile of healthy controls (*n* = 9) and PD patients (*n* = 10), aged 17 to 65 years. Our findings revealed a marked increase in total cholesterol (TC) and triglyceride (TG) along with a decrease in high-density lipoprotein cholesterol (HDL-C) levels, but no significant changes in low-density lipoprotein cholesterol (LDL-C) levels in the serum of PD patients (Fig. 1a and Table S1). To validate these findings, we performed a logistic regression analysis of the characteristic patterns of dyslipidemia (including TC, TG, HDL-C, and LDL-C levels) in relation to the PD pathogenesis based on health survey and oral examination data (Korea National Health and Nutrition Examination Survey (KNHANES) VII (2016 to 2018)) from 11,710 participants, aged 19 to 75 years^21^. According to the logistic regression analysis, dyslipidemia involving hyper-TG and hypo-HDL-C was associated with PD risks (Table 1 and Table S2, S3). To determine the significance of altered lipid profiles in PD patients, we further focused on the pathogenic mechanisms of PD involving abnormal cholesterol metabolism. First, since PD accompanies gingival inflammation, we analyzed the periodontal characteristics of human patient tissues. The expression of pro-inflammatory factors (*MMP1*, *MMP3*, *IL6*, *IL8*, *CCL5*, and *PTGS2*) was upregulated in the gingiva of PD patients compared to that of the healthy controls (Fig. 1b). To investigate the mechanism by which hyperlipidemia affects PD phenotypes, the event related to intracellular accumulation of cholesterol was examined. The use of a cholesterol-specific stain, Filipin, showed an increase in intracellular cholesterol deposits in human inflamed gingiva compared to healthy tissues (Fig. 1c). This finding was further supported by data from PD mouse models (Fig. 1d). Gingival fibroblasts (GF) and periodontal ligament (PDL) cells were the predominant cell types in the periodontal tissue. Stimulation of pro-inflammatory cytokines, IL1β and TNFα, led to upregulated intracellular levels of total cholesterol and free cholesterol in both types of periodontal cells (Fig. 1e). To further explore whether high cholesterol is a potential cause of PD pathogenesis, periodontal cells were treated with cholesterol. Direct cholesterol treatment of primary cultures of human GF and PDL cells led to significantly elevated expression of catabolic factors and inflammatory mediators associated with PD (Fig. 1f and Fig. S1). The hallmark of PD progression is alveolar bone loss which can be evaluated by measuring the distance between the cementum-enamel junction and the alveolar bone crest (CEJ-ABC), as well as bone mineral density (BMD) by μCT or histological analysis^22,23^. To determine whether alveolar bone loss (ABL) is induced by cholesterol, we utilized a ligature-induced PD mouse model and compared the mice fed on a regular diet (RD) vs. those fed on high cholesterol diet (HCD). The results demonstrated a statistically significant increase in ABL and a decrease in BMD in the non-ligature and ligature-induced periodontal tissue of mice fed an HCD compared to the RD group (Fig. 1g). These findings confirmed the association between hypercholesterolemia and PD pathogenesis. Here, we propose a new term, “metabolic PD”, because PD may be a metabolic disease.

**Fig. 1.**
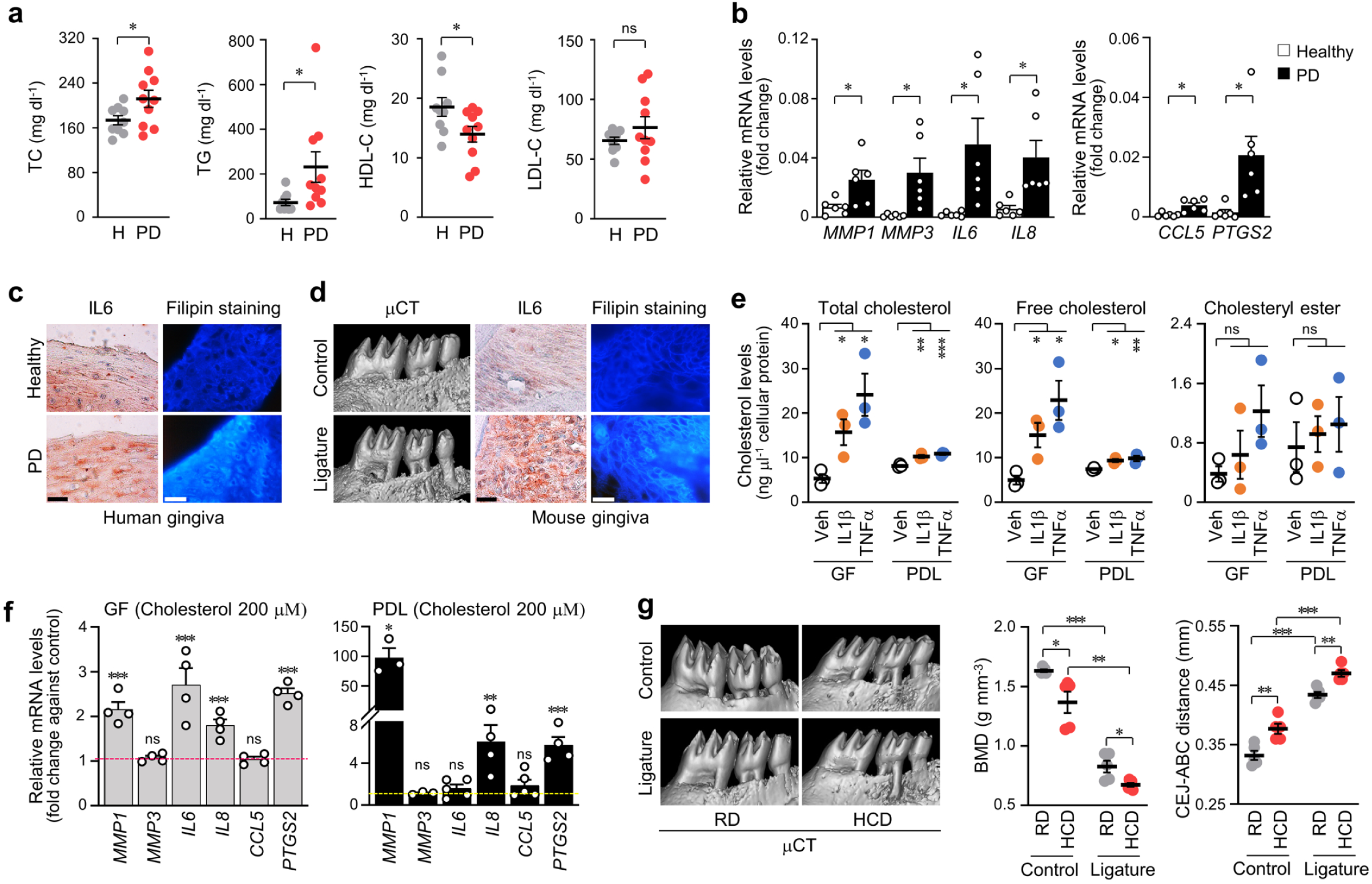
An abnormal cholesterol uptake in the periodontal cells aggravates PD phenotypes. (**a**) Lipid profiling using serum from healthy individuals (non-inflamed, *n* = 9) and patients with chronic PD (inflamed, *n* = 10). (**b)** mRNA levels of the indicated genes involved in PD pathogenesis from non-inflamed (healthy) or inflamed (PD) human gingival tissues (*n* = 6). **(c**, **d)** Representative images of gingival tissues from human patients with PD patients (**c**) and maxilla region of ligature-induced PD mice (**d**) following micro-computed tomography (μCT), IL6 immunostaining, and filipin staining. Scale bar, 25 μm. (**e)** The cellular levels of total cholesterol, free cholesterol, and cholesteryl ester in human GF and PDL cells that were treated with IL1β (2 ng ml^−1^) or TNFα (50 ng ml^−1^)for 24 h (*n* = 3). (**f)** mRNA levels of the indicated genes involved in PD pathogenesis in human GF and PDL cells, treated with cholesterol (200 μM) for 36 h (*n* ≥ 3). (**g)** Representative μCT images and the BMD and CEJ-ABC distance analysis in the maxilla region of ligature-induced PD mice fed an RD or a HCD for 13 weeks (*n* = 5). *n* indicates the number of biologically independent samples or human specimens or mouse number per group. Values are presented as mean ± standard error of the mean (SEM) based on the two-tailed *t*-test (**a**, **b, e, g**) and one-way ANOVA with Tukey’s test (**f**). (\**P* < 0.05, \*\**P* < 0.01, \*\*\**P* < 0.001).

**Table 1.**
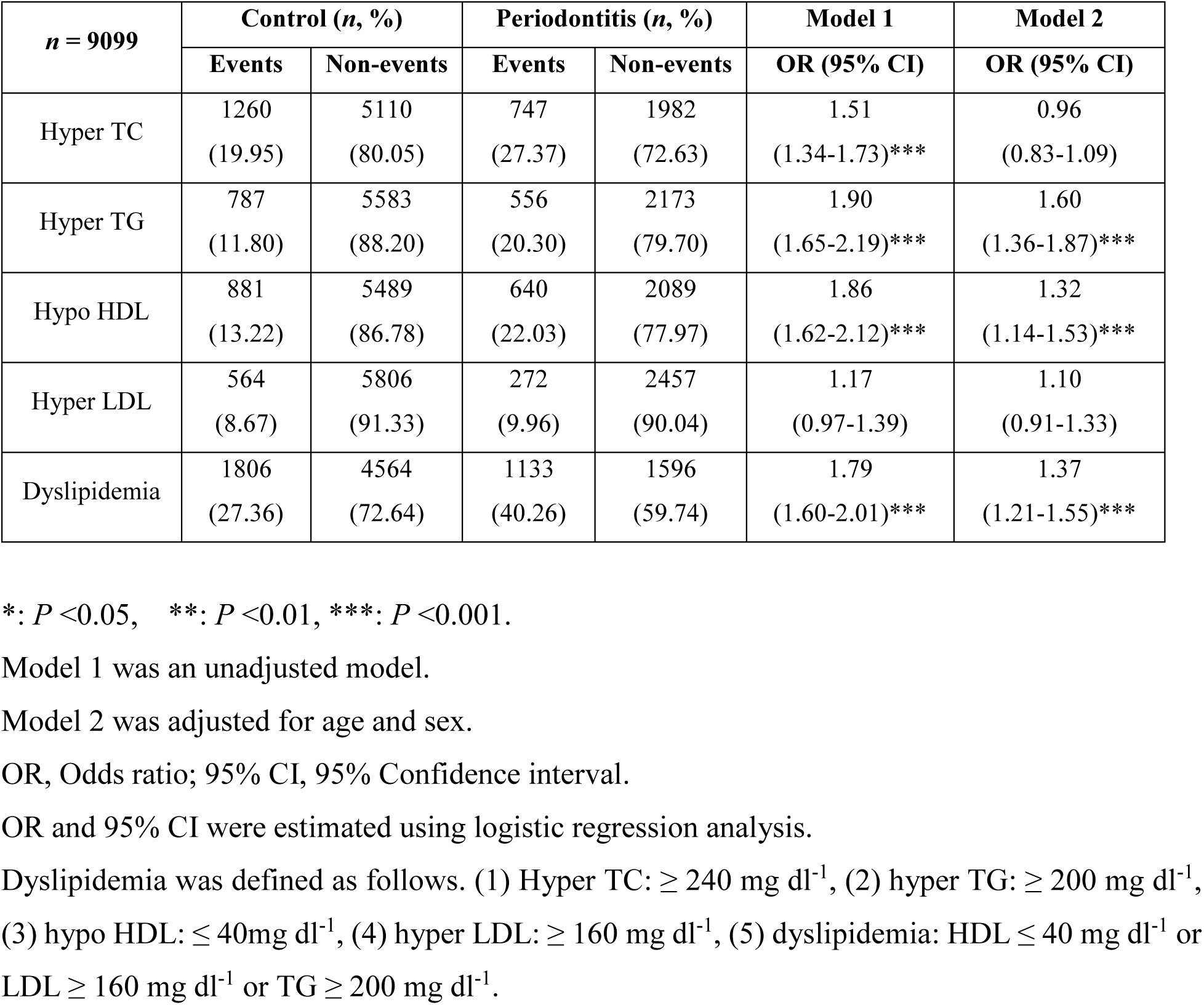
Correlations between hypercholesterolemia and PD: Distribution of dyslipidemia risk factors in human samples with and without periodontitis and results of logistic regression analysis using data from KNHANES VII (2016 to 2018)

### Cholesterol-lowering drugs alleviate metabolic PD

Given the above results, we investigated the effects of clinically available cholesterol-lowering drugs including fenofibrate, atorvastatin, niacin, ezetimibe, and lomitapide, using a PD mouse model (Fig. S2a). Cholesterol-lowering drug treatment was initiated after 4 weeks of RD or HCD and continued for 9 weeks (Fig. S2a). All drugs induced significant recovery of HCD-mediated TC upregulation in serum. Depending on the known mode of action for each drug (Fig. S2b), the group on HCD and fenofibrate showed hyper-HDL-C in serum relative to the group that was fed HCD only. In contrast, those on ezetimibe treatment demonstrated lowered LDL-C levels (Fig. 2a). In male mice with experimental PD, fenofibrate and atorvastatin significantly counteracted the HCD-induced increase in BMD loss and ABL (Fig. 2b, c). Unexpectedly, the individual drug effects on PD phenotypes in female mice differed from the results obtained with male mice (Fig. 2d, e). All examined cholesterol-lowering drugs alleviated the HCD-mediated increase in BMD loss and ABL in experimental female mice, except ezetimibe, which did not affect BMD loss (Fig. 2e). We also examined the effect of lomitapide on alleviating HCD-mediated inflammation and bone loss in mice. The results demonstrated an unexplained drastic decrease in serum TC, HDL-C, and LDL-C (Fig. S3). Next, to verify that the mitigatory effects of cholesterol-lowering drugs on periodontal phenotypes are due to the drug action on serum cholesterol or lipid levels, and not on locally expressed cholesterol or via regulation of other factors (for example, PPAR-α targeting by fenofibrate) by human GF, primary human GF cultures were directly treated with the drugs. The cholesterol-lowering drugs did not demonstrate significant regulatory effects on the IL1β-(Fig. 3a) or TNFα-induced (Fig. 3b) increase in inflammatory gene expression. These findings suggest that the modulatory effects of cholesterol-lowering drugs on periodontal inflammation and the stages following the erosion of alveolar bone may be exerted via the systemic effects of serum cholesterol or lipids. Taken together, our results suggest that hyperlipidemia is a risk factor for the development of metabolic PD and support the potential utility of cholesterol-lowering drugs as a therapeutic strategy.

**Fig. 2.**
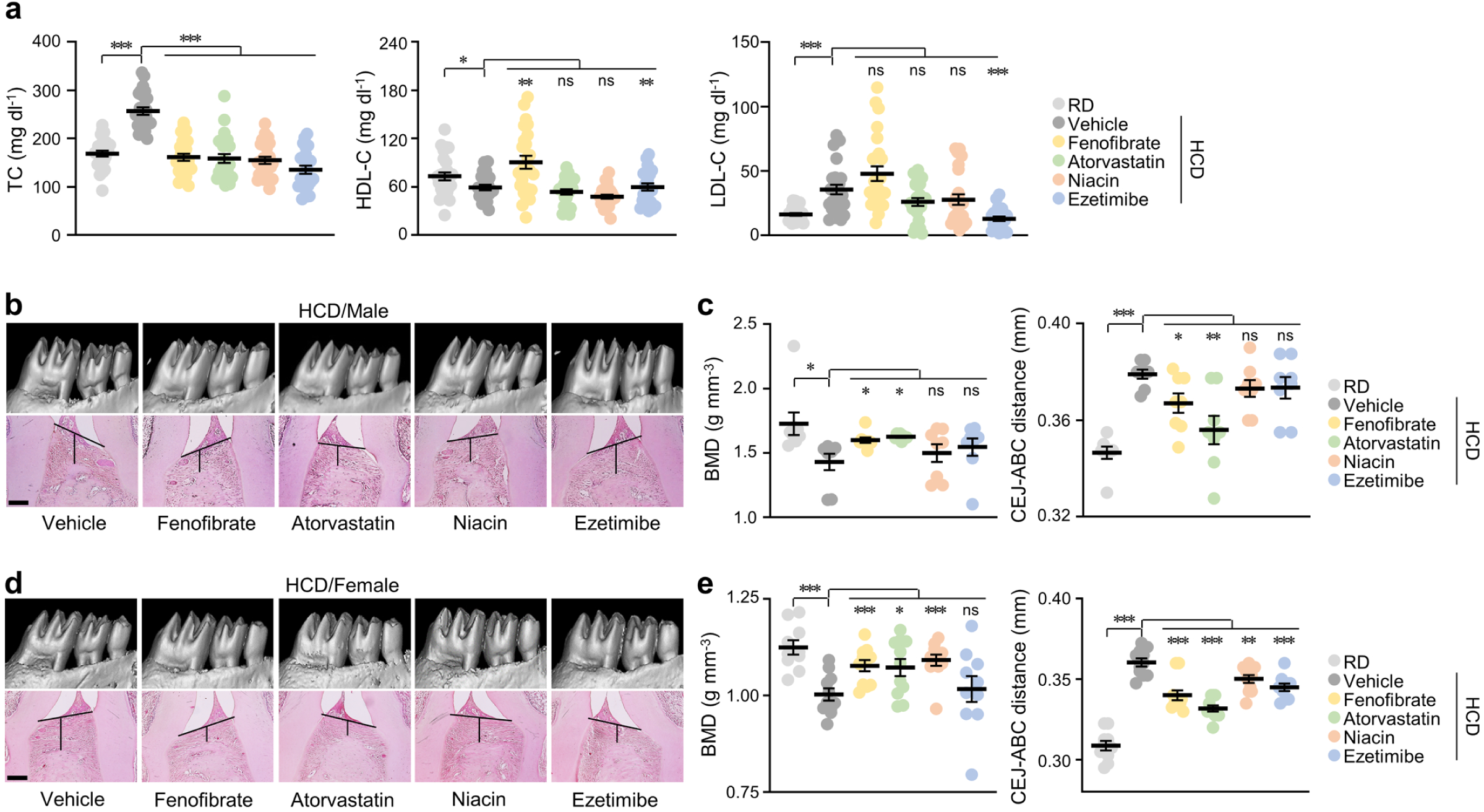
Administration of cholesterol-lowering drugs attenuates HCD-induced metabolic PD. (**a**) TC, HDL-C, and LDL-C levels in serum samples collected from mice that were administered with cholesterol-lowering drugs for 9 weeks and HCD fed for 13 weeks (*n* ≥ 23). (**b**-**e)** Representative μCT images for BMD analysis and H&E staining images for CEJ-ABC distance measurements in the maxilla region of male (**b**, **c**) and female (**d**, **e**) mice administered vehicle (DMSO) or cholesterol-lowering drugs for 9 weeks and HCD feeding for 13 weeks (*n* = 8 (male); *n* ≥ 10 (female)). Scale bar, 100 μm. *n* indicates the number of mice per group. Values are presented as mean ± SEM based on the two-tailed *t*-test (**a**, **c**, **e**). (\**P* < 0.05, \*\**P* < 0.01, \*\*\**P* < 0.001).

**Fig. 3.**
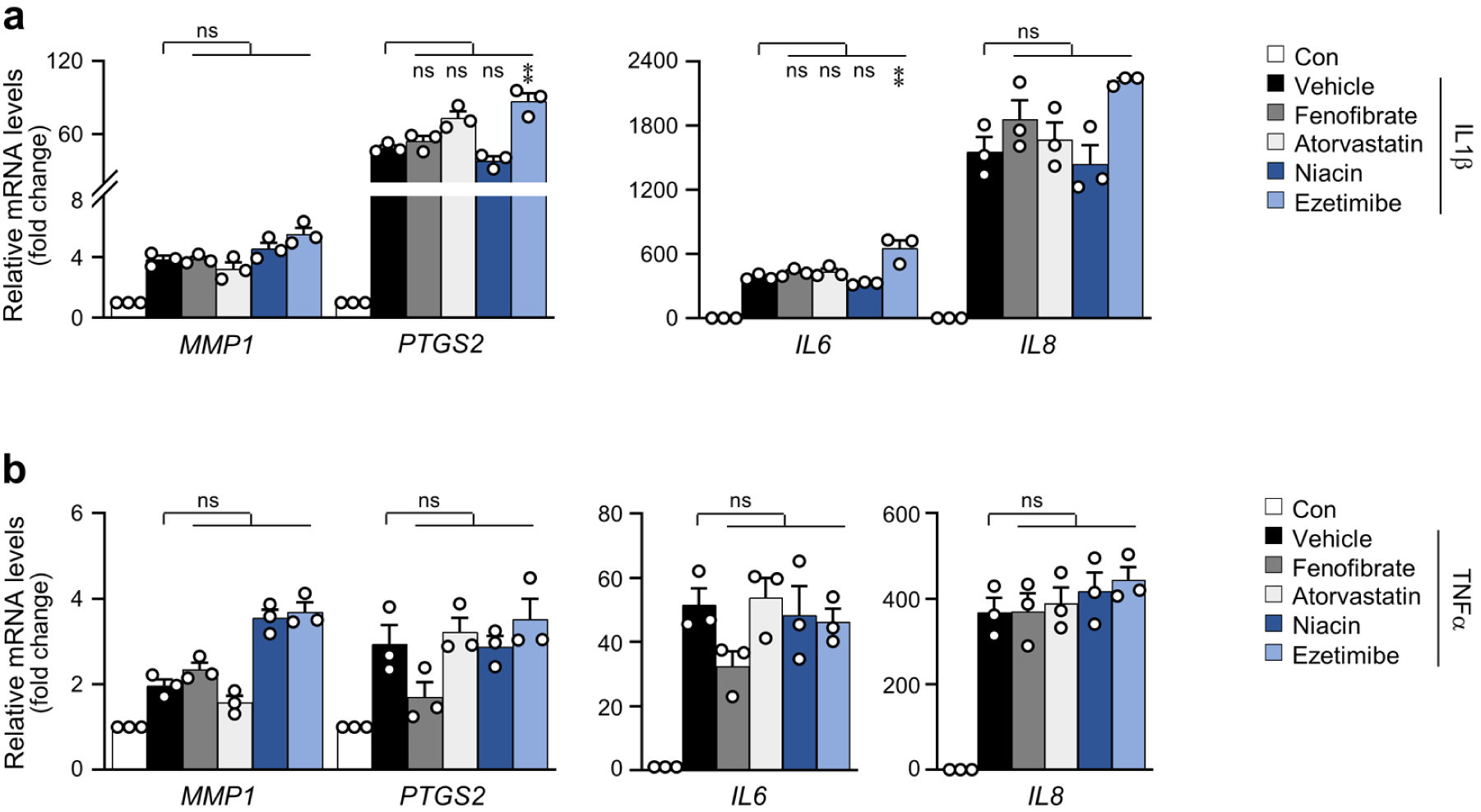
Expression of catabolic factors in human GF treated with cholesterol-lowering drugs. (**a**, **b**) qRT-PCR analysis of *MMP1, PTGS2, IL6, and IL8* in human GF treated with the all cholesterol-lowering drugs (fenofibrate, 25 µM; atorvastatin 1 µM; niacin, 25 µM; ezetimibe, 25 µM) in the presence of IL1β (2 ng ml^−1^) (**a**) or TNFα (50 ng ml^−1^) (**b**) for 24 h. (*n* = 3). DMSO was used as a vehicle. *n* indicates the number of biologically independent samples. Values are presented as mean ± SEM based on one-way ANOVA and Tukey’s test. (\**P* < 0.05, \*\**P* < 0.01, \*\*\**P* < 0.001).

### Cholesterol efflux capacity is inhibited in periodontal inflammation

The protective effect of cholesterol-lowering drugs on PD may be related to the elimination of excess cholesterol. We next examined the modulatory effects of cholesterol efflux mediators on cellular cholesterol levels in PD pathogenesis. Unexpectedly, *ABCA5*, which belongs to the ATP-binding cassette transporter family, was upregulated by both IL1β and TNFα, while significant *ABCA1* upregulation was only detected with IL1β stimulation (Fig. 4a and 4b). Although the upregulation of genes involved in cholesterol efflux occurs in inflamed human GF, it appears that accumulated cholesterol in GF may not be clear due to the downregulation of serum HDL levels in patients with PD (Fig. 1a). Next, we examined whether the stimulation of cholesterol efflux by extracellular HDL treatment could rescue the increased intracellular cholesterol and the subsequent upregulated catabolic gene expression during metabolic PD. With a decrease in intracellular cholesterol level (Fig. 4c), the upregulated expression of catabolic factors including *MMP1*, *IL6*, *IL8*, and *PTGS2*, was significantly suppressed by HDL treatment (Fig. 4d and 4e). These results suggest that cholesterol efflux capacity is inhibited during metabolic PD and that the stimulation of HDL-mediated cholesterol efflux can alleviate metabolic PD pathogenesis.

**Fig. 4.**
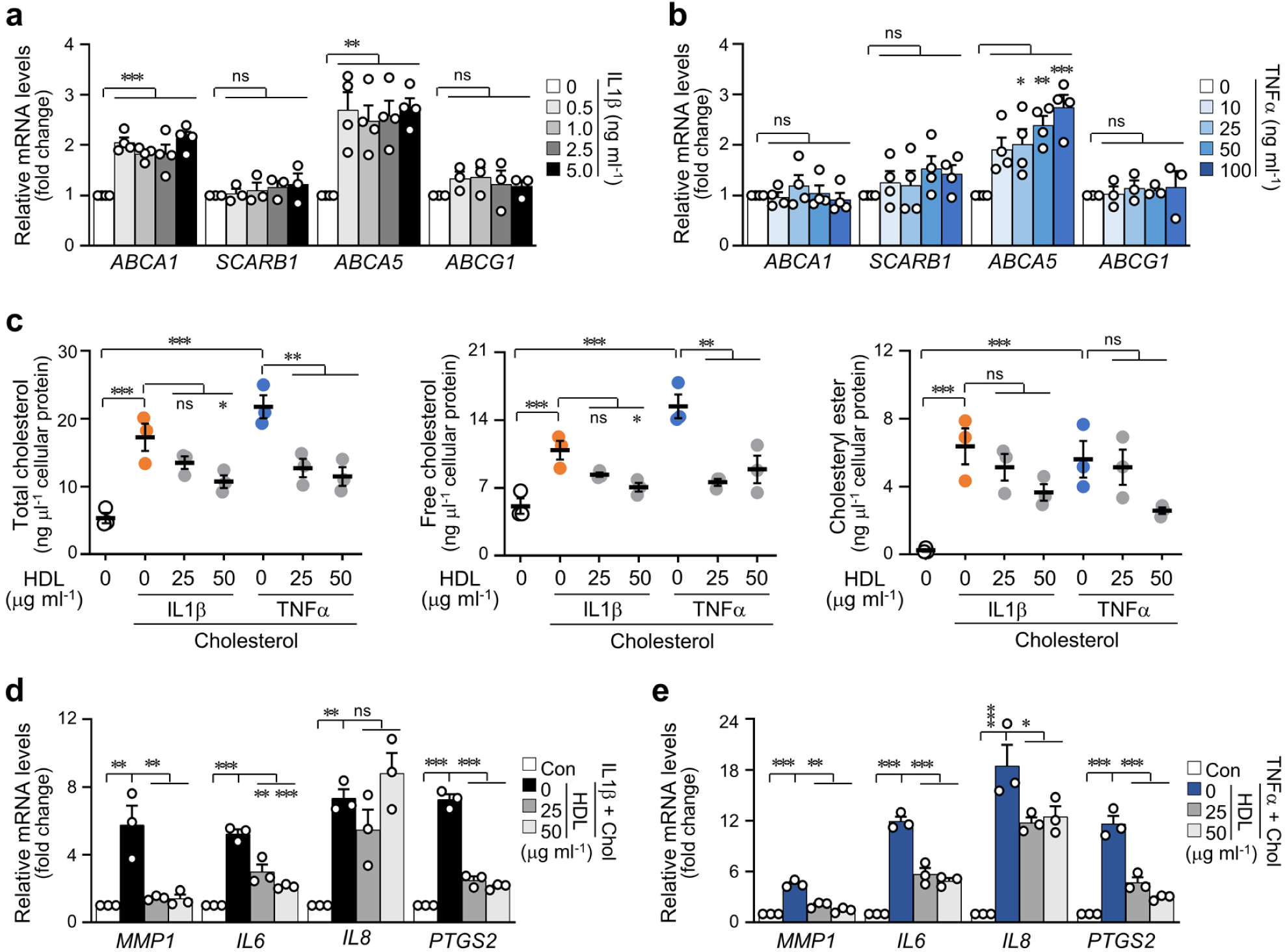
Expression of cholesterol efflux mediators and effect of HDL in human GF treated with pro-inflammatory cytokines. (**a**, **b**) mRNA levels of cholesterol efflux mediators (*ABCA1*, *SCARB1*, *ABCA5*, *ABCG1*) in human GF treated with IL1β (**a**) or TNFα (**b**) for 24 h (*n* ≥ 3). (**c)** Cellular levels of total cholesterol, free cholesterol, and cholesteryl ester in cholesterol (25 μM) pre-loaded human GF treated with HDL in the presence of IL1β (2 ng ml^−1^) or TNFα (50 ng ml^−1^) for 12 h (*n* = 3). (**d**, **e)** mRNA levels of the indicated genes involved in PD in cholesterol (25 μM) pre-loaded human GF treated with HDL in the presence of IL1β (2 ng ml^−1^) or TNFα (50 ng ml^−1^) for 12 h (*n* = 3). *n* indicates the number of biologically independent samples. Values are presented as mean ± SEM based on one-way ANOVA and Tukey’s test. (\**P* < 0.05, \*\**P* < 0.01, \*\*\**P* < 0.001).

### Acceleration of APOA1BP-mediated cholesterol efflux prevents PD exacerbation

HDL particles comprise a lipid core (cholesteryl esters, triglycerides, and free cholesterol) and apolipoproteins (APOA1, APOA2, APOE), as well as antioxidants and enzymes related to plasma lipid metabolism including paraoxonase 1 (PON1), hepatic lipase (LIPC), endothelial lipase (LIPG), phospholipid transfer protein (PLTP), cholesteryl ester transfer protein (CETP), and lecithin-cholesterol acyltransferase (LCAT)^24,25^. The biological functionality of HDL is principally influenced by the composition and regulator. Therefore, we examined for defects in the regulators of cholesterol efflux or HDL components in inflamed human GF induced by IL1β (Fig. 5a) or TNFα (Fig. 5b). Unlike other efflux-related genes with no change at the transcript level, our results showed the downregulation of *APOA1BP* (a booster of cholesterol efflux) by IL1β but not TNFα^26^ (Fig. 5c and 5d). The suppression of the APOA1BP protein expression in inflamed human (Fig. 5e) and mouse PD tissues (Fig. 5f) was confirmed. Next, we validated the protective role of APOA1BP in PD pathogenesis. The overexpression of *APOA1BP* induced a reduction in the expression of inflammatory genes and catabolic factors by IL1β (Fig. 6a) and cholesterol (Fig. 6b), but not by TNFα (Fig. S4a). The inhibitory effects of *APOA1BP* overexpression on IL1β-or cholesterol-mediated increase in all cellular cholesterol types were significant (Fig. 6c and Fig. S4b), while the TNFα-mediated upregulation of cholesterol was not altered (Fig. S4c). The role of APOA1BP in PD pathogenesis was further confirmed *in vivo* by intra-gingival injection of Ad-*Apoa1bp* in mice. Stimulation of cholesterol efflux via *APOA1BP* overexpression inhibited the ligature-induced PD phenotypes (Fig. 6d and 6e). Collectively, the results suggest that the downregulation of APOA1BP in periodontal gingiva is a key event in the blockage of cholesterol efflux. The results also support the therapeutic potential of drugs that activate APOA1BP-induced cholesterol efflux for the management of metabolic PD.

**Fig. 5.**
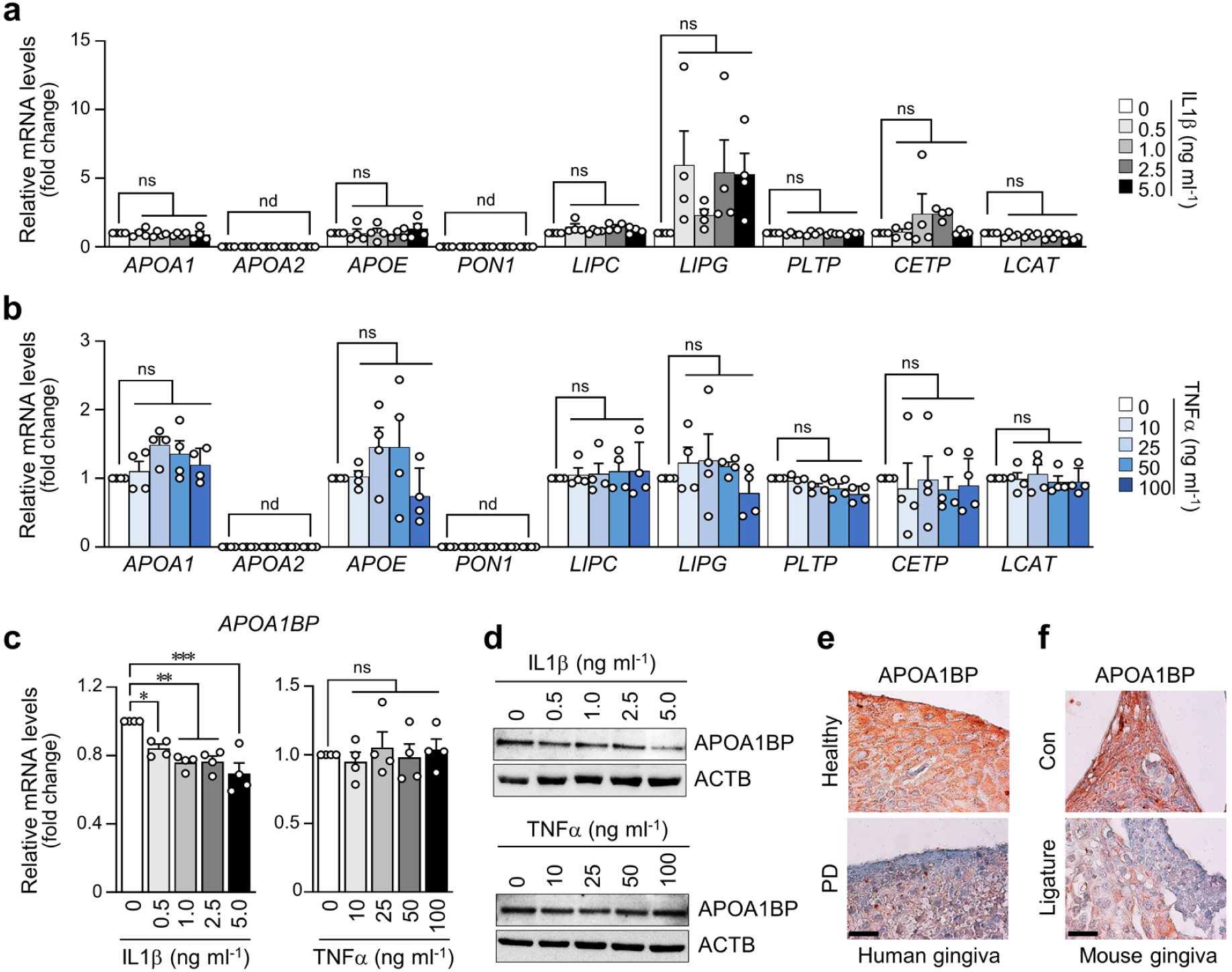
Upregulation of APOA1BP in human GF treated with pro-inflammatory cytokines and in periodontal tissues. (**a**, **b**) mRNA levels of HDL-related genes (HDL component: *APOA1*, *APOA2*, *APOE*, *PON1*; HDL interaction: *LIPC*, *LIPG*, *PLTP*, *CETP*, *LCAT*) in human GF treated with IL1β (**a**) or TNFα (**b**) for 24 h (*n* = 4). **c** *APOA1BP* mRNA levels in human GF treated with IL1β or TNFα for 24 h (*n* = 4). **d** Protein expression of APOA1BP in human GF treated with IL1β or TNFα for 24 h. (**e, f)** Representative APOA1BP immunostaining images in gingiva from healthy, and PD patients (**e**) and ligature-induced PD mice (**f**). Scale bar, 25 μm. *n* indicates the number of biologically independent samples. Values are presented as mean ± SEM based on one-way ANOVA and Tukey’s test. (\**P* < 0.05, \*\**P* < 0.01, \*\*\**P* < 0.001).

**Fig. 6.**
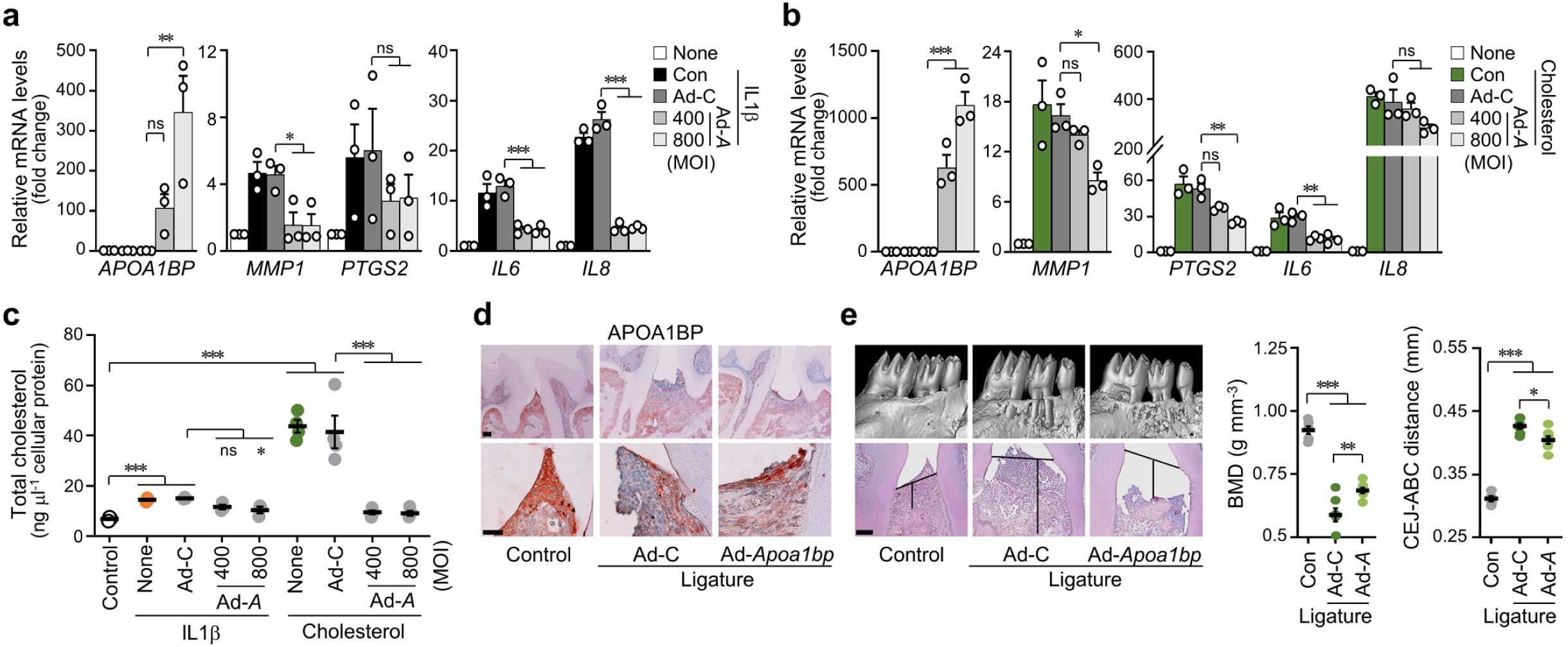
APOA1BP-mediated cholesterol efflux alleviates PD progression. (**a**, **b**) *APOA1BP*, *MMP1*, *PTGS2, IL6, and IL8* mRNA levels in human GF infected with Ad-*APOA1BP* (Ad-*A*) in the presence of IL1β (2 ng ml^−1^) (**a**) or cholesterol (200 μM) (**b**) for 48 h (*n* = 3). **c** Total cholesterol levels in human GF infected with Ad-C or Ad-*APOA1BP* (Ad-*A*) in the presence of IL1β (2 ng ml^−1^) or cholesterol (200 μM) for 24 h (*n* = 4). (**d**, **e)** Representative APOA1BP immunostaining, μCT, and H&E staining images for BMD analysis and CEJ-ABC distance in the maxilla region of ligature-induced PD mice injected intra-gingivally with Ad-C and Ad-*Apoa1bp* (1 × 10^9^ PFU per 3 μl) (*n* = 7). Scale bar, 100 μm (immunostaining image – above). Scale bar, 25 μm (immunostaining image – below). Scale bar, 100 μm (H&E staining image). *n* indicates the number of biologically independent samples or the number of mice per group. Values are presented as mean ± standard error of the mean (SEM) based on the two-tailed *t*-test (**e**) and one-way ANOVA with Tukey’s test (**a**, **b**, **c**). (\**P* < 0.05, \*\**P* < 0.01, \*\*\**P* < 0.001).

## DISCUSSION

PD is primarily caused by pathogenic microorganisms, but its severity is known to be positively associated with systemic diseases^27–29^. Studies on both human and animal models suggest a bi-directional association between PD and metabolic disorders^27,30,31^. In particular, whether hyperlipidemia is a risk factor for PD is yet to be established. This research question was based on the findings that hyperlipidemia is one of the major risk factors for cardiovascular diseases including atherosclerotic vascular disorder. Our results confirmed that systemic hypercholesterolemia is a potent causative factor in PD development, and the alterations in gene expression patterns, induced by the inflammatory responses in periodontal cells cooperatively promote PD pathogenesis. Furthermore, we suggest that the stimulation of HDL-mediated cholesterol efflux results in the effective alleviation of PD phenotypes.

Data from a range of meta-analyses suggest that PD is positively associated with lipid profiles^32^. Since both PD and dyslipidemia are affected by age, genetic background, eating habits, nutritional status, and disease status, we conducted a logistic regression analysis using Korean population data, which confirmed a strong positive correlation between PD and dyslipidemia (Table 1, S2, and S3). Hypo-HDL-C and hyper-TG status obtained from human serum samples were consistent with the clinical data (Fig. 1a), although the observed levels of TC were inconsistent. A multitude of variables may have ramifications in the association between PD and lipid profiles, including study populations and subjects, which are yet to be determined. The relationship between PD and dyslipidemia was stronger with age (Table S2), thus supporting an aging-dyslipidemia-PD axis. To maintain cholesterol homeostasis in cells, excess cellular cholesterol is released by extrahepatic cells and carried to the liver for reuse in bile synthesis or excretion in the process of reverse cholesterol transport^33,34^. HDL works as the primary acceptor for the released cholesterol, and apoA-I, the key apoprotein in HDL, predominantly mediates cholesterol efflux^35,36^. Thus, hypo-HDL-C in patients with PD can disrupt cholesterol efflux in periodontal cells, thereby, potentially leading to the accumulation of intracellular cholesterol. However, the upregulation of efflux-associated genes under inflammatory conditions did not correlate with the accumulation of intracellular cholesterol in our experiments. To solve this paradox, we examined the functional defects of the efflux-associated factors. Our results revealed the downregulation of APOA1BP in PD pathogenesis. Notably, APOA1BP modulation was observed with IL1β signaling, but not TNFα signaling. This finding supports the hypothesis that APOA1BP is involved in the upregulation of efflux-associated genes. In contrast, ABCA5 was upregulated under both IL1β– and TNFα-signaling. These findings suggest that ABCA1 is responsible for APOA1BP-mediated cholesterol efflux. However, future studies are required for further validation.

The protective efficacies of clinically available cholesterol-lowering drugs (fenofibrate, statin, niacin, ezetimibe, and lomitapide)^37^ with their different modes of action on PD pathogenesis were further compared in mice. Cholesterol is obtained as a form of chylomicron from the diet and endogenous biosynthesis^38^. Fenofibrate, an agonist of PPARα, activates lipolysis and eliminates TG-rich particles from the plasma. Fenofibrate has been utilized to treat specific populations with hyper-TG and hypo-HDL-C^39^. Statins target the rate-limiting step in de novo synthesis of cholesterol that is catalyzed by 3-hydroxy-3methylglutaryl coenzyme A reductase (HMGCR)^40^. Niacin, a non-competitive inhibitor of diacylglycerol acyltransferase2 (DGAT2), suppresses TG synthesis in hepatocytes^41^. Ezetimibe inhibits the absorption of dietary cholesterol by blocking Niemann-Pick C1-like 1 (NPC1L1), a cholesterol transporter expressed in enterocytes^42,43^. Lomitapide inhibits the microsomal triglyceride transfer protein (MTP), which is necessary for VLDL assembly and secretion in the liver^44^ and is also located in the endoplasmic reticulum of enterocytes^45,46^. However, the molecular targets of these drugs may be more complex, leading to diverse effects. In our study, all the examined drugs induced a decrease in HCD-induced hyper-TC but each exerted distinct effects on the HDL-C and LDL-C levels. HCD-induced hyper-LDL-C was reduced by ezetimibe and lomitapide, while HCD-induced hypo-HDL-C was increased by fenofibrate. Atorvastatin and fenofibrate exerted significant protective effects on PD phenotypes, periodontal inflammation, and subsequent alveolar bone loss in both male and female mice. Consistently, several earlier studies suggest that statins administered to patients with chronic PD demonstrated additional anti-inflammatory effects^47–49^. Our data not only validated the efficacy of atorvastatin against PD in mice but also demonstrated the regulation of catabolic gene expression through cholesterol-associated metabolic pathways. Indeed, fenofibrate exerted a significant effect on periodontal inflammation in both male and female mice. The results demonstrated by fenofibrate indicate that *Apoa1bp*-mediated cholesterol efflux alleviates metabolic PD pathogenesis, and thus, may offer an effective therapeutic strategy for metabolic PD that target periodontal cells.

Certain cholesterol-lowering drugs exerted sex-dependent effects in mice (Fig. 2). Moreover, logistic regression analysis using data from KNHANES VII results indicated a stronger relationship between PD and dyslipidemia in male compared to female patients (Table S3). Interestingly, previous studies reported that the rate of PD in men (∼55%) is higher than in women (∼38%)^9,50^, implying a potential gender bias in PD pathogenesis. These sex-dependent results may be attributable to the effects of hormones, or some unknown factors. In this study, we showed for the first time that the efficacy of cholesterol-lowering drugs in mice was due to changes in cholesterol metabolism in the periodontal cells. The specific signaling pathways of each drug involved in the inhibition of metabolic PD phenotypes were not examined in this study and required further investigation. Based on the preliminary findings, we propose that combined treatment with currently available cholesterol-lowering medications may present an effective approach to mitigate the pathogenesis of metabolic PD. In addition, further experiments to determine the exact role of sex in PD pathogenesis are warranted.

The results of this study provided direct evidence that cholesterol homeostasis is disrupted in PD conditions. Moreover, an increased intracellular cholesterol level in periodontal cells is one of the causative factors of PD development, thus, a rebalance of cholesterol homeostasis by HDL treatment may help in alleviating periodontal inflammation. Furthermore, our findings also demonstrate that the prevention and alleviation of metabolic PD are necessary for oral health and the maintenance of normal physiological processes. Our study supports the utility of combined treatment with clinically available cholesterol-lowering drugs as an effective approach for the management of PD.

## MATERIALS AND METHODS

### Logistic regression analysis of data from the Korea National Health and Nutrition Examination Survey

To elucidate the correlation between periodontitis and dyslipidemia, raw data from the Korea National Health and Nutrition Examination Survey (KNHANES) VII (2016 to 2018) conducted by the Korea Disease Control and Prevention Agency and Prevention and Ministry of Health and Welfare were analyzed. KNHANES consisted of three surveys: health, screening, and nutritional surveys. In this study, data from the health and oral examination surveys of participants, aged 19 to 75 years, were collected and analyzed. Of the 11,948 participants, 9099 were included in the analysis, after excluding those with missing periodontal information. Data were obtained following the approval of the Institutional Review Board of the Korea Disease Control and Prevention Agency (2018-01-03-P-A). All participants provided signed informed consent prior to participation. Oral examinations were performed by trained dentists according to KNHANES guidelines. For the diagnosis of periodontitis, the Community Periodontal Index (CPI) recommended by WHO was used. The evaluation criteria of CPI are classified into five categories as follows. Code 0 (sound periodontal tissue), code 1 (bleeding periodontal tissue), code 2 (calculus formation in periodontal tissue), code 3 (periodontal pocket of 4-5 mm), and code 4 (deep periodontal pocket ≥ 6 mm). Results were defined as follows. Codes 0-2 were classified as “no periodontitis” and codes 3 and 4 as “periodontitis”. For the evaluation of dyslipidemia, the TC, TG, and HDL levels were measured directly from blood tests, and low-density lipoprotein (LDL) values were obtained using the Friedewald formula^51^. To determine the prevalence of lipid factors, criteria of the Committee of Clinical Practice Guidelines of the Korean Society of Lipid and Atherosclerosis were used^52^. The prevalence of dyslipidemia was defined as follows. (1) Hyper TC: ≥ 240 mg dl^−1^, (2) hyper TG: ≥ 200 mg dl^−1^, (3) hypo HDL: ≤ 40mg dl^−1^, (4) hyper LDL: ≥ 160 mg dl^−1^, (5) dyslipidemia: HDL ≤ 40 mg dl^−1^ or LDL ≥ 160 mg dl^−1^ or TG ≥ 200 mg dl^−1^.

### Human subjects

We gathered human gingival tissues from twelve individuals (mean age = 49.17 ± 12.40 years; range = 34 to 65 years old) during tooth extraction. We obtained six non-inflamed gingival tissue samples from healthy persons and six inflamed gingival tissue samples from patients with chronic periodontitis. The Institutional Review Board of Chonnam National University Dental Hospital (Gwangju, Republic of Korea) approved this study. The patients provided informed consent after all the protocols had been clearly explained. The human gingiva tissue samples were promptly preserved in liquid nitrogen and placed at –80°C for the next analysis. Human blood was obtained from 9 healthy subjects without periodontitis (mean age = 32.56 ± 15.39 years; range = 17 to 65 years old) and 10 patients with periodontitis (mean age = 52.20 ± 6.76 years; range = 39 to 59 years old). For serum analysis, the collected blood was centrifuged at 7000 rpm for 10 minutes at 4°C, and supernatant fractions were obtained. Sera were stored at –80°C until analysis.

### Mice

All animals were reared in pathogen-free barrier facilities at the Chonnam National University. This research was approved by the Animal Care and Ethics Committee of Chonnam National University.

### Administration of a high-cholesterol diet and cholesterol-lowering drugs to mice

C57BL/6 male and female mice (5 weeks old) were fed AIN-76A (CLS bio, Gyeonggi-do, Republic of Korea) as the RD or a modified AIN-76A diet supplemented with 2% cholesterol as the HCD (CLS bio). After 4 weeks of HCD feeding, the mice were given HCD containing five different cholesterol-lowering drugs (fenofibrate 150 mg kg**^-^**^1^ day**^-^**^1^, atorvastatin 6 mg kg**^-^**^1^ day**^-^**^1^, niacin 360 mg kg**^-^**^1^ day**^-^**^1^, ezetimibe 7 mg kg**^-^**^1^ day**^-^**^1^ or lomitapide 4 mg kg**^-^**^1^ day**^-^**^1^, CLS bio) for 9 weeks. After this period, mice were sacrificed, and the maxilla was harvested at 13 weeks for μCT and histological analyses.

### Establishment of experimental PD in the mouse model

A silk ligature 5-0 (Ailee, SK521, Busan, Republic of Korea) was placed around the maxillary right second molar of 10-week-old male mice, and the left side was used as the control^53^. A group of mice with ligature-induced PD were injected intra-gingivally with Ad-*Apoa1bp* (1 × 10^9^ plaque-forming units (PFU) in a total volume of 3 μl) once per day for 7 days. The control mice received the same ligature surgery but were administered an intra-gingival injection with empty adenovirus (Ad-C). Mice were sacrificed after 9 days of ligation and periodontal tissue samples were collected^53^. All adenoviruses were provided by Vector Biolabs.

### Primary cell culture

Healthy human gingiva and ligament tissues at the middle third of the roots were obtained during tooth extraction^54^. The Institutional Review Board of Chonnam National University Hospital approved the protocol and informed written consent was obtained from all participants. The specimens were minced into small pieces in phosphate-buffered saline solution (PBS) and directly cultured in Dulbecco’s modified Eagle’s medium (DMEM, Gibco, 12800-017, Massachusetts, USA) containing 10% fetal bovine serum (FBS), 1% penicillin and 1% streptomycin. Human GF and PDL cells between the 4^th^ and 9^th^ passages were used. Cells were treated with cholesterol (3β-hydroxy-5-cholestene, Sigma-Aldrich, C-3045; cholesterol–methyl-β-cyclodextrin, Sigma-Aldrich, C-4951, Missouri, USA), IL1β (GenScript, Z02922-10, New Jersey, USA) or TNFα (GenScript, Z02774-20). Human GF was treated with five cholesterol-lowering drugs (fenofibrate, orb322488; atorvastatin, orb134747; niacin, orb321956; ezetimibe, orb134389; lomitapide, orb251455, Biorbyt, Cambridge, UK) in the presence of IL1β (2 ng ml^−1^) or TNFα (50 ng ml^−1^). Human GF was infected with *APOA1BP*-overexpressing adenovirus in the presence of IL1β (2 ng ml^−1^), TNFα (50 ng ml^−1^), or cholesterol (200 μM). Cells were pretreated with cholesterol (200 μM) for 12 hours and washed twice with PBS. The high-density lipoproteins (HDL; Lee Biosolutions, 361-12, Missouri, USA) were subsequently treated with IL1β (2 ng ml^−1^) or TNFα (50 ng ml^−1^) for 12 hours.

### Micro-computed tomography (μCT)

All maxillae were fixed in 10% neutral buffered formalin (NBF) for 24 hours and measured using the μCT scanner (1172 SkyScan, Belgium). The X-ray generator was operated at 49 kV and 200 μA with a 0.7 mm aluminum filter at 11 μm resolution. Linear and volume analysis of alveolar bone in the first and the second molar was assessed based on 2-D and 3-D outcomes, using CTAn (SkyScan) and Mimics 14.0 (Materialise, Belgium).

### Histological analysis of gingival tissue

Human gingival tissues, with or without inflammation, were fixed in 10% NBF for 24 hours. The mice maxillae were placed in 0.5 M ethylenediamine tetra-acetic acid (EDTA, pH 8.0) for 2 weeks for decalcification. All the samples were embedded in paraffin and sectioned at 5 μm thickness for hematoxylin and eosin (H&E) and immunohistochemical staining. Immunohistochemistry to detect the expression of APOA1BP in the gingiva was performed using primary antibodies rabbit anti-APOA1BP (Biorbyt, orb155691; 1:150). The stained sections were analyzed under a Zeiss Axio Scope A1 microscope.

### Cholesterol assay and analysis of the lipid profiles

Total and free cellular cholesterol concentrations in human GF and PDL cells were determined by a total cholesterol assay kit (Biomax, BM-CHO-100, Gyeonggi-do, Republic of Korea). The levels of TC, TG, HDL-C, and LDL-C were measured in sera collected from human patients with PD or HCD mice, who were administered cholesterol-lowering diets using a total cholesterol assay kit (Biomax), triglyceride assay kit (Biomax, BM-TGR-100) and HDL, LDL/VLDL assay kit (Biomax, BM-CDL-100), respectively. All analyses were performed according to the manufacturer’s protocols.

### Filipin staining

Human and mouse samples were transferred as frozen tissues and sectioned at 10 μm. Filipin (250 mg ml^−1^; Sigma-Aldrich, F9765) staining was carried out on slide-mounted sections following the manufacturer’s instructions. Images were viewed under a Zeiss Axio Scope A1 microscope connected to a fluorescence component, and analyzed by Image J software (National Institutes of Health, v.1.51a).

### Real-time quantitative PCR analysis

Total RNA was extracted from cells or human gingival tissues using TRIzol reagent and converted to cDNA. Real-time RT–PCR was executed on an iCycler (Bio-Rad Laboratories, California, USA). The primers are listed in Table S4. All qRT-PCR reactions were duplicated, and all samples were normalized using the ΔΔCt method.

### Immunoblot analysis

Cells were gently lysed in RIPA buffer, including a protease inhibitor cocktail (Roche, 04906845001, Basel, Switzerland) and a phosphatase inhibitor cocktail (Roche, 11697498001) for 30 min followed by centrifugation and the supernatant was collected as a protein extract. The blots were probed with mouse anti-ACTB (Sigma-Aldrich, A3854) and rabbit anti-APOA1BP (Biorbyt). Membranes were washed with 0.1% Tween in Tris-buffered saline, followed by incubation with horseradish peroxidase-conjugated secondary antibodies (anti-rabbit IgG; Sigma-Aldrich, A6154). Protein expression was detected using ImageSaver6 software via an EZ capture MG system (ATTO).

### Statistical analysis

Since KNHANES is a complex sample survey, a complex sample data analysis was conducted. To investigate the effects of lipids on periodontitis, complex sample multiple logistic regression analysis was performed. Data were analyzed using SAS 9.4 software (SAS Institute, inc., Cary, North Carolina, USA). The statistical significance was set to *P* < 0.05. All experiments were conducted at least three times. Values were reported as mean ± SEM. Statistical analyses were performed using SPSS 25 and GraphPad Prism 8 software. A two-tailed *t-*test was applied to compare differences between two groups, and one-way ANOVA, followed by Tukey’s post hoc test (multi-comparison), was used for the comparison of means among more than two independent groups. The *n* value represented the number of independent experiments or mice. A *P* value of less than 0.05 was considered to indicate significance.

## Acknowledgments

This work was supported by National Research Foundation of Korea (NRF) grants, funded by the Korean government (MSIT) (2019R1A5A2027521 and 2021R1A2C3005727), and a Korean Fund for Regenerative Medicine (KFRM) grant funded by the Korean government (the Ministry of Science and ICT, the Ministry of Health & Welfare) (22A0104L1).

## Author Contributions

T.-T.T. and G.L. designed, performed, and analyzed the experiments. K.-H.C. performed the logistic regression analysis. S.Y.L. and K.H.P. assisted with the experimental design and analyses. M.-S.K., J.R., O.-S.K., and H.-P.L. provided human serum and tissues. Y.H.H. and J.-T.K. analyzed and evaluated mouse PD models. T.-T.T., G.L., Y.H.H., and J.-H.R. wrote the manuscript. J.-H.R. conceived, planned, and oversaw the study. All authors provided critical revision of the manuscript and approved the final version for submission.

