## Supplemental Information for "Acceleration of HDL-mediated cellular cholesterol efflux alleviates periodontitis"

Supplemental Figure 1 to 4

Supplemental Table 1 to 4

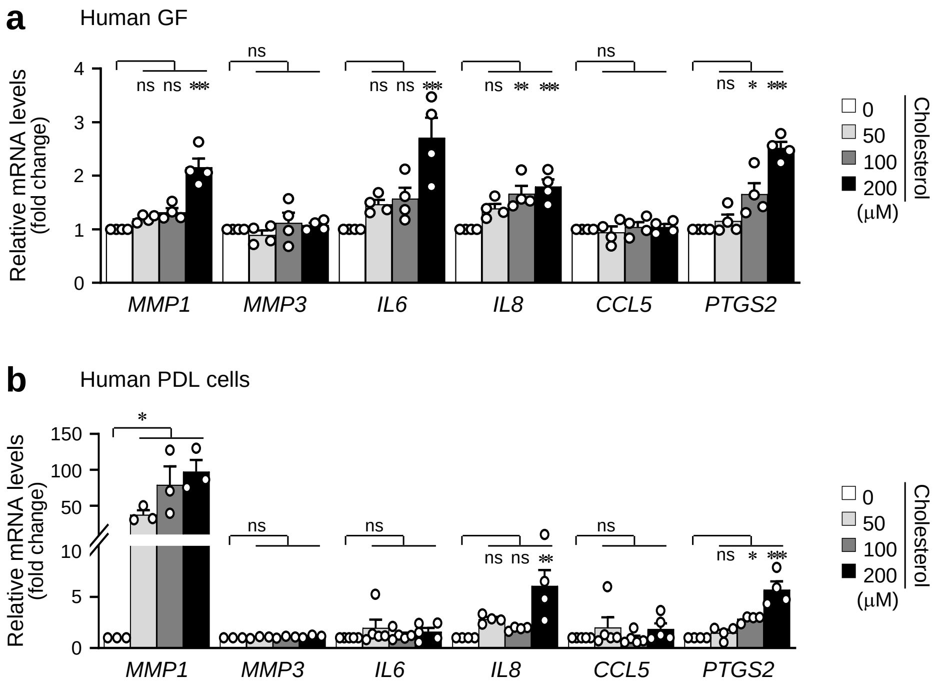

**Fig. S1. Expression of catabolic factors in human GF and PDL cells treated with cholesterol (a, b)** qRT-PCR analysis of *MMP1, MMP3, IL6, IL8, CCL5,* and *PTGS2* in human GF (**a**) and PDL cells (**b**) treated with the indicated amount of cholesterol for 24 h. ($n \geq 3$). $n$ indicates the number of biologically independent samples. Values are presented as mean ± SEM based on one-way ANOVA and Tukey's test. (*$P < 0.05$, **$P < 0.01$, ***$P < 0.001$).

a

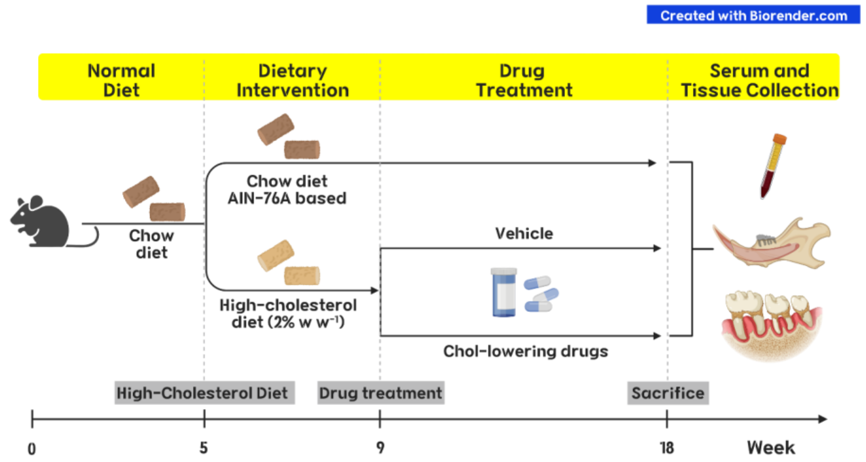

b

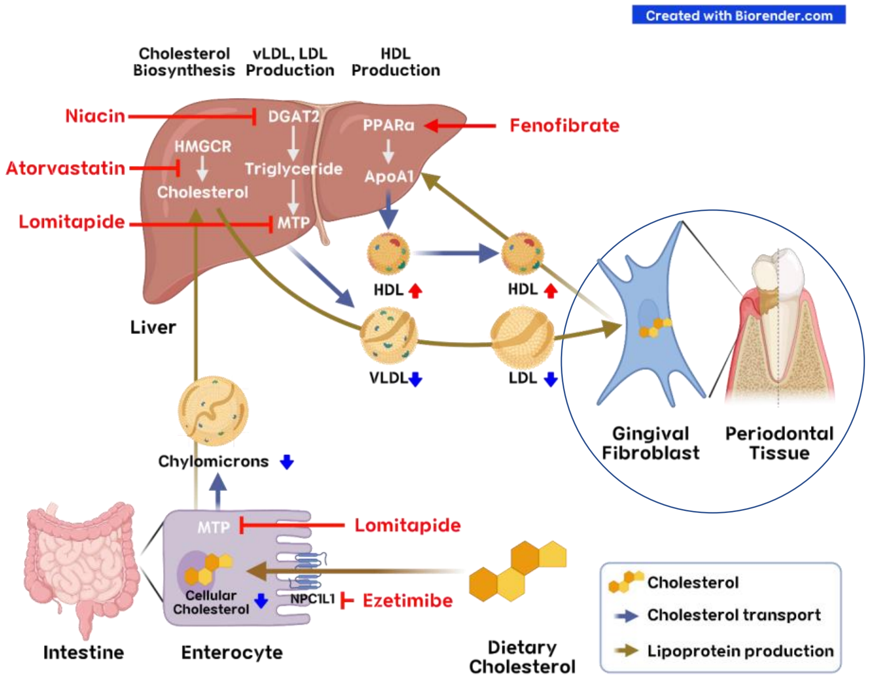

**Fig. S2. Schematic diagram of the mechanisms of action of the cholesterol-lowering drugs and experimental procedures**

**(a)** The schematic diagram of experimental procedures of male and female mice fed RD or HCD with the administration of cholesterol-lowering drugs for 13 weeks, including fenofibrate (150 mg kg$^{-1}$ day$^{-1}$), atorvastatin (6 mg kg$^{-1}$ day$^{-1}$), niacin (360 mg kg$^{-1}$ day$^{-1}$), ezetimibe (7 mg kg$^{-1}$ day$^{-1}$), and lomitapide (4 mg kg$^{-1}$ day$^{-1}$). **(b)** The schematic diagram of the mechanisms of action of cholesterol-lowering drugs regulating cholesterol transport into periodontal tissues. These schematic diagrams were created with BioRender.com.

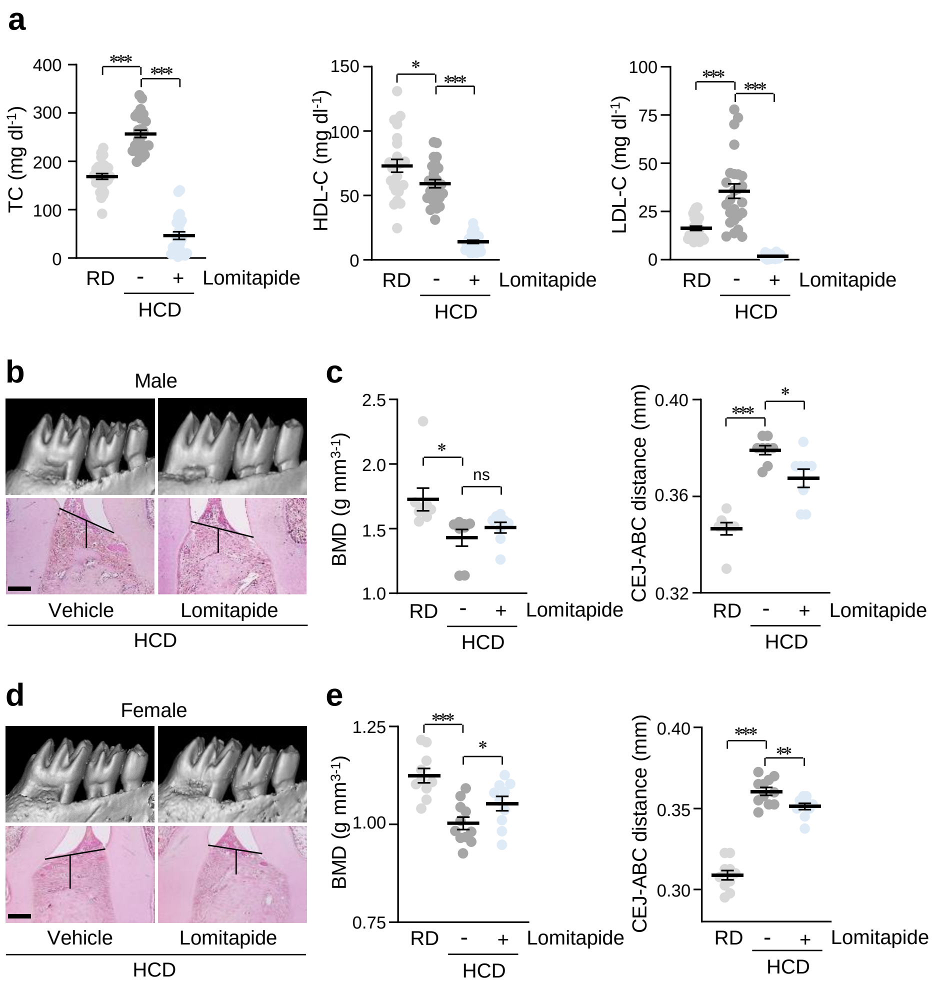

**Fig. S3. The effect of lomitapide administration on PD pathogenesis**

**(a)** TC, HDL-C, and LDL-C levels in serum collected from mice administered with lomitapide for 9 weeks during HCD feeding for 13 weeks ($n \geq 25$). **(b-e)** Representative µCT images with BMD analysis, and H&E staining images with CEJ-ABC distance measurements in the maxilla region of the male (**b**, **c**) and female (**d**, **e**) mice administered with vehicle or lomitapide for 9 weeks during HCD feeding for 13 weeks ($n = 8$ (male); $n \geq 10$ (female)). Scale bar, 100 µm. *n* indicates the number of mice per group. Values are presented as mean ± SEM. A two-tailed *t*-test was used (*$P < 0.05$, **$P < 0.01$, ***$P < 0.001$).

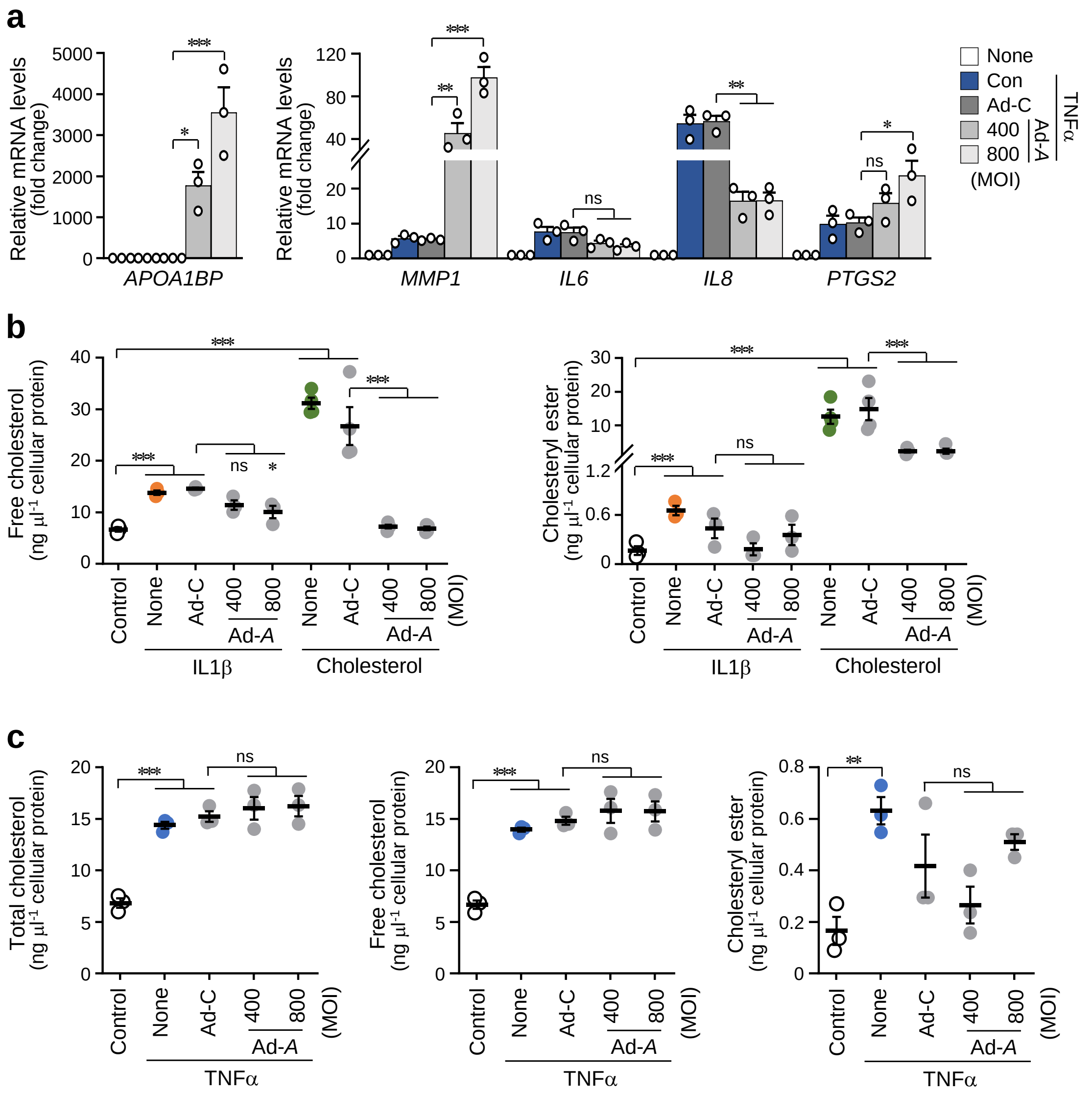

**Fig. S4. Effect of ectopic overexpression of APOA1BP on catabolic gene expression in human GF treated with TNFα**

**(a)** *APOA1BP*, *MMP1*, *IL6*, *IL8,* and *PTGS2* mRNA levels in human GF infected with Ad-*APOA1BP* (Ad-*A*) in the presence of TNFα (50 ng $ml^{-1}$) for 48 h ($n$ = 3). **(b)** Free cholesterol (left) and cholesteryl ester (right) levels in human GF infected with Ad-*APOA1BP* in the presence of IL1β (2 ng $ml^{-1}$) or cholesterol (200 µM) for 24 h ($n$ = 4).**(c)** Cellular cholesterol levels in human GF infected with Ad-*APOA1BP* in the presence of TNFα (50 ng $ml^{-1}$) for 48 h ($n$ = 3). $n$ indicates the number of biologically independent samples. Values are presented as mean ± SEM based on one-way ANOVA and Tukey's test (*$P < 0.05$, **$P < 0.01$, ***$P < 0.001$).

**Table S1. Lipid profiling of serum samples collected from healthy individuals (non-inflamed, *n* = 9) and patients with chronic PD (inflamed, *n* = 10)**

| **Human Serum** | **Healthy (*n* = 9)** | **PD (*n* = 10)** | ***P*-value** |
|---|---|---|---|
| Sex (M, F) | 6, 3 | 7, 3 | |
| Age (years) | 32.56 ± 15.39 | 52.20 ± 6.76 | |
| Total Cholesterol (mg $dl^{-1}$) | 173.722 ± 8.282 | 212.121 ± 15.509 | 0.0497* |
| Triglyceride (mg $dl^{-1}$) | 73.421 ± 13.796 | 231.80 ± 68.377 | 0.0472* |
| HDL-C (mg $dl^{-1}$) | 18.537 ± 1.583 | 13.963 ± 1.329 | 0.0396* |
| LDL-C (mg $dl^{-1}$) | 65.519 ± 3.088 | 76.546 ± 9.231 | 0.2814 |

**Table S2. Distribution of dyslipidemia components in young (19~39) and aged (40~75) subjects with and without periodontitis and result of logistic regression analysis**

| **Age: 19~39** | **Control (*n*, %)** | | **Periodontitis (*n*, %)** | | **Model 1** | **Model 2** |
|---|---|---|---|---|---|---|
| | **Events** | **Non-events** | **Events** | **Non-events** | **OR (95% CI)** | **OR (95% CI)** |
| Hyper TC | 197 (7.84) | 2318 (92.16) | 33 (12.55) | 219 (87.45) | 1.69 (1.06-2.68)* | 1.18 (0.74-1.87) |
| Hyper TG | 246 (9.52) | 2269 (90.48) | 57 (20.74) | 195 (79.26) | 2.49 (1.67-3.70)*** | 1.60 (1.09-2.34)* |
| Hypo HDL | 280 (10.77) | 2235 (89.23) | 51 (18.08) | 201 (81.92) | 1.83 (1.28-2.62)*** | 1.23 (0.86-1.76) |
| Hyper LDL | 147 (5.79) | 2368 (94.21) | 25 (8.54) | 227 (91.46) | 1.5 2(0.90-2.57) | 1.14 (0.69-1.90) |
| Dyslipidemia | 551 (21.15) | 1964 (78.85) | 99 (34.74) | 153 (65.26) | 1.98 (1.41-2.80)*** | 1.29 (0.92-1.80) |
| **Age: 40~75** | **Control (*n*, %)** | | **Periodontitis (*n*, %)** | | **Model 1** | **Model 2** |
| | **Events** | **Non-events** | **Events** | **Non-events** | **OR (95% CI)** | **OR (95% CI)** |
| Hyper TC | 1063 (27.78) | 2792 (72.22) | 714 (28.88) | 1763 (71.12) | 1.06 (0.93-1.20) | 0.95 (0.83-1.09) |
| Hyper TG | 541 (13.27) | 3314 (86.73) | 499 (20.25) | 1978 (79.75) | 1.66 (1.42-1.94)*** | 1.51 (1.28-1.79)*** |
| Hypo HDL | 601 (14.80) | 3254 (85.20) | 589 (22.43) | 1888 (77.57) | 1.66 (1.44-1.93)*** | 1.33 (1.13-1.55)*** |
| Hyper LDL | 417 (10.52) | 3438 (89.48) | 247 (10.10) | 2230 (89.90) | 0.96 (0.79-1.16) | 1.05 (0.85-1.28) |
| Dyslipidemia | 1255 (31.37) | 2600 (68.63) | 1034 (40.82) | 1443 (59.18) | 1.51 (1.33-1.71)*** | 1.33 (1.17-1.52)*** |

*: $P < 0.05$, **: $P < 0.01$, ***: $P < 0.001$.

Model 1 was an unadjusted model.

Model 2 was adjusted for sex.

OR, Odds ratio; 95% CI, 95% Confidence interval.

OR and 95% CI were estimated using logistic regression analysis.

Dyslipidemia was defined as follows. (1) Hyper TC: ≥ 240 mg $dl^{-1}$, (2) hyper TG: ≥ 200 mg $dl^{-1}$, (3) hypo HDL: ≤ 40mg $dl^{-1}$, (4) hyper LDL: ≥ 160 mg $dl^{-1}$, (5) dyslipidemia: HDL ≤ 40 mg $dl^{-1}$ or LDL ≥ 160 mg $dl^{-1}$ or TG ≥ 200 mg $dl^{-1}$.

**Table S3. Distribution of dyslipidemia components in male and female subjects with and without periodontitis and result from the logistic regression analysis**

| **Men (*n* = 3853)** | **Control (*n*, %)** | | **Periodontitis (*n*, %)** | | **Model 1** | **Model 2** |
|---|---|---|---|---|---|---|
| | **Events** | **Non-events** | **Events** | **Non-events** | **OR (95% CI)** | **OR (95% CI)** |
| Hyper TC | 442 (18.03) | 1965 (81.97) | 331 (22.21) | 1115 (77.79) | 1.30 (1.08-1.56)** | 0.96 (0.79-1.17) |
| Hyper TG | 458 (18.48) | 1949 (81.52) | 389 (26.47) | 1057 (73.53) | 1.59 (1.33-1.89)*** | 1.74 (1.45-2.10)*** |
| Hypo HDL | 547 (22.28) | 1860 (77.72) | 460 (30.00) | 986 (70.00) | 1.50 (1.25-1.79)*** | 1.31 (1.08-1.60)** |
| Hyper LDL | 191 (7.63) | 2216 (92.37) | 113 (7.88) | 1333 (92.12) | 1.04 (0.76-1.41) | 1.14 (0.83-1.56) |
| Dyslipidemia | 930 (37.69) | 1477 (62.31) | 718 (48.12) | 728 (51.88) | 1.53 (1.31-1.80)*** | 1.47 (1.24-1.75)*** |

| **Women (*n* = 5246)** | **Control (*n*, %)** | | **Periodontitis (*n*, %)** | | **Model 1** | **Model 2** |
|---|---|---|---|---|---|---|
| | **Events** | **Non-events** | **Events** | **Non-events** | **OR (95% CI)** | **OR (95% CI)** |
| Hyper TC | 1818 (21.06) | 3145 (78.94) | 416 (32.78) | 867 (67.22) | 1.83 (1.56-2.14)*** | 0.92 (0.77-1.10) |
| Hyper TG | 329 (7.96) | 3634 (92.04) | 167 (13.82) | 1116 (86.18) | 1.85 (1.45-2.37)*** | 1.37 (1.05-1.79)* |
| Hypo HDL | 334 (8.01) | 3629 (91.99) | 180 (13.66) | 1103 (86.34) | 1.82 (1.46-2.26)*** | 1.32 (1.05-1.67)* |
| Hyper LDL | 373 (9.26) | 3590 (90.74) | 159 (12.14) | 1124 (87.86) | 1.35 (1.08-1.70)** | 1.05 (0.82-1.35) |
| Dyslipidemia | 876 (21.43) | 3087 (7.57) | 415 (32.01) | 868 (67.99) | 1.73 (1.48-2.02)*** | 1.25 (1.06-1.49)** |

*: $P$ <0.05, **: $P$ <0.01, ***: $P$ <0.001.

Model 1 was an unadjusted model.

Model 2 was adjusted for age.

OR, Odds ratio; 95% CI, 95% Confidence interval.

OR and 95% CI were estimated using logistic regression analysis.

Dyslipidemia was defined as follows. (1) Hyper TC: ≥ 240 mg $dl^{-1}$, (2) hyper TG: ≥ 200 mg $dl^{-1}$, (3) hypo HDL: ≤ 40mg $dl^{-1}$, (4) hyper LDL: ≥ 160 mg $dl^{-1}$, (5) dyslipidemia: HDL ≤ 40 mg $dl^{-1}$ or LDL ≥ 160 mg $dl^{-1}$ or TG ≥ 200 mg $dl^{-1}$.

**Table S4. PCR primer sequences and PCR conditions**

| Gene | Strand | Primer sequences | Size (bp) | AT (°C) | Origin |
|---|---|---|---|---|---|
| *ABCA1* | S | 5’-GCTCTCAGACCTGGGCATTT-3’ | 260 | 60 | Human |
| | AS | 5’-CCTTTGCCATCCATCCCACT-3’ | | | |
| *ABCA5* | S | 5’-TTCTATGTCCTCTTGGCTGTCTATC-3’ | 252 | 60 | Human |
| | AS | 5’-TTCACCCTTCTTTCTGTATGTCTTC-3’ | | | |
| *ABCG1* | S | 5’-GCGTGCCTGGGTGATGAGAAATAATG-3’ | 272 | 60 | Human |
| | AS | 5’-ATGATGACAATGCTTGAGCCCTGTAG-3’ | | | |
| *APOA1BP* | S | 5’-TGTGCTCGACACCTCAAACT-3’ | 307 | 60 | Human |
| | AS | 5’-TTTCCCTTCTCCACGTCCCA-3’ | | | |
| *APOA1* | S | 5’-CTAAACCTAAAGCTCCTTGACAACTG-3’ | 213 | 60 | Human |
| | AS | 5’-CATCTCCTCCTGCCACTTCTTCTG-3’ | | | |
| *APOA2* | S | 5’-GGAGAGCCTGGTTTCTCAGTACTTC-3’ | 200 | 60 | Human |
| | AS | 5’-GCTGTGTTCCAAGTTCCACGAAATAG-3’ | | | |
| *APOAE* | S | 5’-GCTGCGTTGCTGGTCACATTCCTG-3’ | 286 | 60 | Human |
| | AS | 5’-TCAGTTGTTCCTCCAGTTCCGATTTG-3’ | | | |
| *CCL5* | S | 5’-GGATCAAGACAGCACGTGGA-3’ | 421 | 60 | Human |
| | AS | 5’-CGGGTGGGGTAGGATAGTGA-3’ | | | |
| *CETP* | S | 5’-CAGCTGTTCACAAATTTCATCTCCTT-3’ | 252 | 60 | Human |
| | AS | 5’-GAAGATTTCCTGGTTGGTGTTGAAG-3’ | | | |
| *GAPDH* | S | 5’-GGTGAAGGTCGGAGTCAACG-3’ | 327 | 60 | Human |
| | AS | 5’-CAAATGAGCCCCAGCCTTCT-3’ | | | |
| *IL6* | S | 5’-AGGCTGGACTGCAGGAACTCCTTAAAG-3’ | 421 | 60 | Human |
| | AS | 5’-CCCTGAGAAAGGAGACATGTAACAAGAG-3’ | | | |
| *IL8* | S | 5’-CTCTCTTGGCAGCCTTCCTGATTTC-3’ | 254 | 60 | Human |
| | AS | 5’-AAACTTCTCCACAACCCTCTGCAC-3’ | | | |
| *LCAT* | S | 5’-TTCATTTCCACACCCAGCTTCAACTA-3’ | 245 | 60 | Human |
| | AS | 5’-TCATCACCATCCTCATAGAGCACAC-3’ | | | |
| *LIPC* | S | 5’-AACAACTTTTACCATGTCACTACTCG-3’ | 321 | 60 | Human |
| | AS | 5’-AGTAGTAGGTCATCTGTGTTTTCTGA-3’ | | | |
| *LIPG* | S | 5’-TTCGGCTTGAGCATTGGTATTCAGA-3’ | 314 | 60 | Human |
| | AS | 5’-CTCTTGTTCCTCATTTTCTTGGCATT-3’ | | | |
| *MMP1* | S | 5’-GGAGGGGATGCTCATTTTGATG-3’ | 541 | 60 | Human |
| | AS | 5’-TAGGGAAGCCAAAGGAGCTGT-3’ | | | |
| *MMP3* | S | 5’-AATCCTACTGTTGCTGTGCGTG-3’ | 238 | 60 | Human |
| | AS | 5’-CAGAGTGTCGGAGTCCAGCTTC-3’ | | | |

| | | | | | |
|---|---|---|---|---|---|
| *PLTP* | S | 5’-CAGGAGGAAGAGCGGATGGTGTATGTG-3’ | 288 | 60 | Human |
| | AS | 5’-GGCAATGGTGACGCTAGCAGTGACAG-3’ | | | |
| *PON1* | S | 5’-AGAGCTTCAACCCCAACAGTC-3’ | 388 | 60 | Human |
| | AS | 5’-ATACGACCACGCTAAACCCA-3’ | | | |
| *PTGS2* | S | 5’-AATCCTTGCTGTTCCCACCCATG-3’ | 329 | 60 | Human |
| | AS | 5’-AAGGGAGTCGGGCAATCATCAGG-3’ | | | |
| *SCARB1* | S | 5’-AGCAAAACTGTAGGGTCCTGA-3’ | 499 | 60 | Human |
| | AS | 5’-GCACTGAGTCCCCACTGAAT-3’ | | | |
| *Apoa1bp* | S | 5’-CCCCCACGTCTATGTCCAAG-3’ | 499 | 60 | Mouse |
| | AS | 5’-AGCAGGTGGTACAAAGCGAC-3’ | | | |

AT, annealing temperature; S, sense; AS, antisense.
